# IgG sialylation occurs via the FcRn-mediated recycling pathway in endothelial cells

**DOI:** 10.1101/2023.06.30.547255

**Authors:** Leandre M. Glendenning, Kalob M. Reynero, Emily N. Kukan, Megan D. Long, Brian A. Cobb

**Affiliations:** Department of Pathology, Case Western Reserve University; Cleveland, OH, 44124, USA

## Abstract

IgG is a key mediator of immune responses throughout the human body, and the structure of the conserved glycan on the Fc region has been identified as a key inflammatory switch regulating its downstream effects. In particular, the absence of terminal sialic acid has been shown to increase the affinity of IgG for activating Fc receptors, cascading the inflammatory response in a variety of diseases and conditions. Previously, we have shown that IgG sialylation is mediated by B cell-extrinsic processes. Here, we show that the FcRn-mediated recycling pathway within endothelial cells is a critical modulator of IgG sialylation. Building a deeper understanding of how IgG sialylation is regulated will drive the development of novel therapeutics which dynamically tune IgG functionality *in vivo*.

**One-Sentence Summary:** Endothelial cells remodel IgG glycans within the FcRn-mediated recycling pathway.

## Main Text

Antibodies, particularly the subtype IgG, play a key role in mediating the immune response through a variety of effector functions. While the specificity of IgG is determined by the sequence and conformation of the Fab domain, its effector functions are largely determined by the Fc domain and its ability to bind to activating or inhibitory Fc receptors (FcR) on immune cells. A crucial component of its ability to bind to these receptors is the composition of a conserved glycan at Asn297 (*1–4*). Specifically, the presence of a terminal sialic acid on this glycan shifts the FcR affinity balance towards the inhibitory Fc receptor FcγRIIB, dampening the downstream immune response. Decreases in IgG sialylation, conversely, lead to an enhanced inflammatory response due to an increase in affinity for activating Fc receptors (*5–8*). The role of IgG sialylation in inflammation has been characterized across a wide variety of diseases, ranging from autoimmune conditions such as rheumatoid arthritis (RA) to infections such as tuberculosis and HIV (*8–17*). However, despite the clear importance of IgG sialylation in the pathology of these diseases, it is unclear how this process is regulated.

Previously, we created a murine B cell-specific knockout of the siayltransferase ST6Gal1, the enzyme responsible for adding terminal sialic acid in an α-2,6 linkage (*18*). This siayltransferase has been localized to the *trans-*Golgi network (*19*), and it is believed that terminal sialic acids are added as a secreted protein passes through the secretory pathway. Thus, we hypothesized that knocking out ST6Gal1 in the B cells of this animal would yield a lack of sialic acids on all of the surface and secreted proteins on these cells. As expected, ST6Gal1 appeared to be necessary for B cell surface sialylation; however, we found that B cell-localized ST6Gal was unnecessary for sialylation of circulating IgG. In follow-up studies, we found that B cells are inefficient at IgG sialylation due to a lack of access to the enzyme ST6Gal1 during secretion (*20*). These data show that IgG glycan composition is modulated following its secretion from the B cell (*20*). We have since determined that IgG sialylation also does not occur within the bloodstream (*21*) or via endocytosis to granules in platelets (*22*) despite its apparent access to ST6Gal1 within these compartments.

IgG lifetime is dramatically enhanced compared to most plasma-localized glycoproteins due to the action of the recycling Fcγ receptor FcRn (*23–26*). Endothelial cells lining the blood vessels pinocytose plasma components and shuttle those macromolecules into endo/lysosomes that are subsequently acidified. The low pH not only activates local proteases to degrade ingested proteins, but also promotes IgG binding to FcRn, which selectively rescues the antibody from degradation by re-releasing it back into the circulatory environment. Here, we report that IgG glycans are sialylated within the FcRn-mediated recycling pathway of endothelial cells (*27*), but not FcRn-expressing macrophages (*23, 28*). This discovery paves the way to begin understanding the regulatory features which drive disease-mediated changes in IgG function via glycan remodeling.

### FcRn knockout animals have decreased plasma IgG sialylation

It has been long understood that IgG in the plasma of FcRn knockout mice (FcRnKO) has a significantly shorter half-life than wild-type mice, presumably due to the lack of FcRn-mediated protection from the intracellular lysosomal pathway (*23, 27, 27, 29*). As IgG half-life also correlates with IgG sialylation (*30*), we hypothesized that IgG isolated from the plasma of FcRnKO animals would also be deficient in IgG sialylation. Thus, we examined the composition of the glycan on isolated IgG through a panel of plant lectins, such as Sambucus nigra (SNA), which binds specifically to α2,6-linked terminal sialic acids. We found that the sialylation status of IgG isolated from the plasma of FcRnKO animals was significantly lower than that of wild-type animals (Fig. 1A). Plasma IgG sialylation in this animal was not as low as in animals completely lacking the enzyme ST6Gal1, suggesting that FcRn may not be strictly necessary, but it dramatically increases the efficiency of IgG sialylation.

**Fig. 1.**
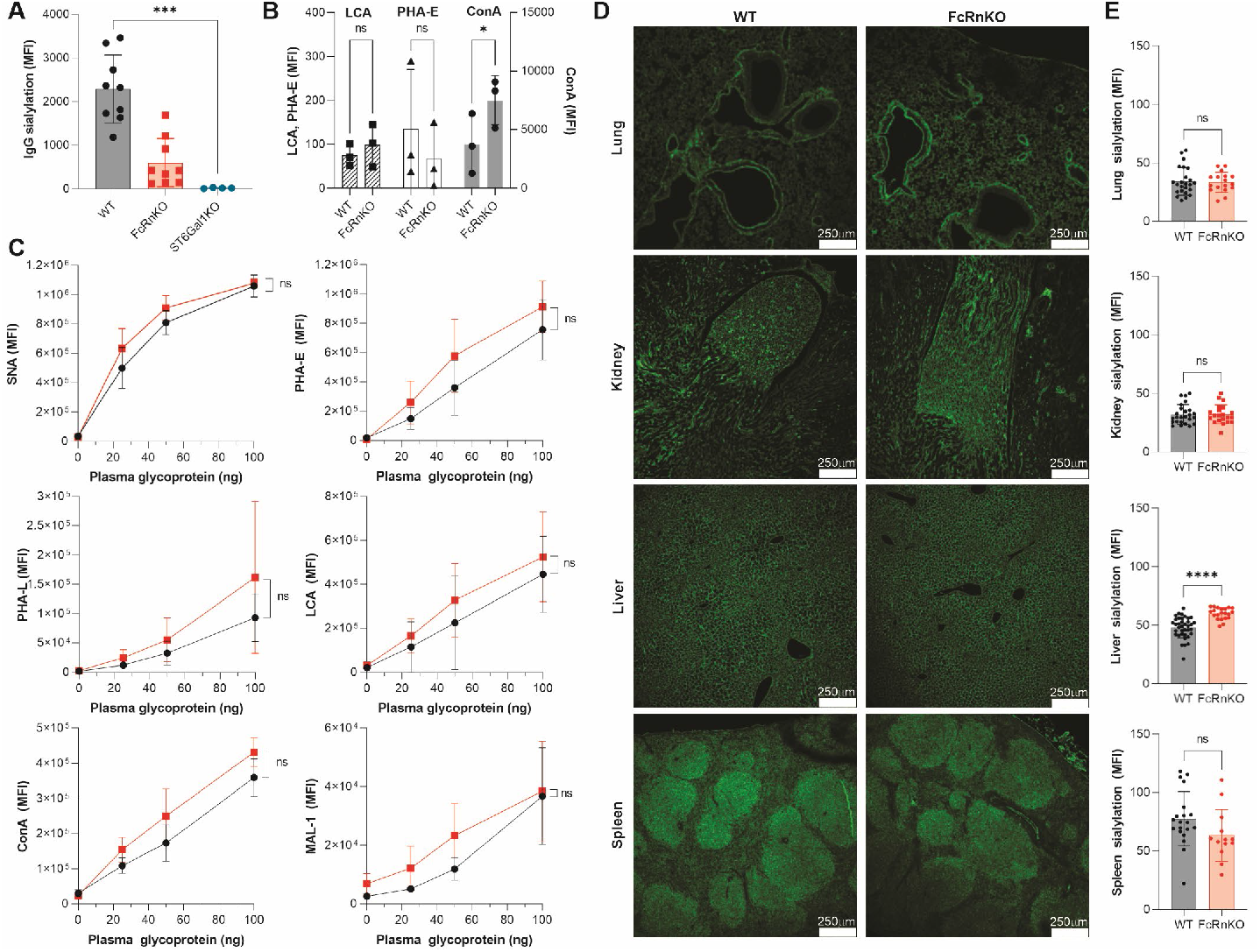
FcRn knockout animals display decreased plasma IgG sialylation. IgG was isolated via high performance liquid chromatography (HPLC) from the plasma of n = 3 mice. IgG was then coated on an ELISA plate, and glycan structure was assessed via lectin ELISA. Sialylation as measured by SNA signal was significantly reduced in FcRnKO and germline ST6Gal1KO animals (**A**), while mannosylation (ConA), core fucosylation (LCA), and bisecting GlcNAc (PHA-E) were not significantly changed between wild type and FcRnKO animals (**B**). Total plasma glycoproteins were assayed via the same technique, and no significant changes were noted in sialylation (SNA), β1,6 branching (PHA-L), mannosylation (ConA), bisecting GlcNAc (PHA-E), core fucosylation (LCA), or α2,3 sialylation (MAL-1) (**C**). No overt pathology was noted in tissue sections of FcRnKO animals (**D**) nor significant differences in tissue sialylation (**E**).

We also interrogated other structural features of the IgG glycan in these animals, such as mannosylation via concanavalin A (ConA) binding, core fucosylation via lens culinaris agglutinin (LCA), and bisecting GlcNAc via phaseolus vulgaris erythroagglutinin (PHA-E). Unlike the difference in SNA signal between the FcRnKO and WT animals, no significant differences in the signal from any of these lectins was found (Fig. 1B).

We next investigated whether these changes in glycan structure are specific to IgG from the circulation of these animals by performing a lectin panel on the bulk plasma glycoproteins from the plasma of FcRnKO and WT animals. There were no differences in sialylation (as measured by SNA), bisecting GlcNAc (PHA-E), β1,6 branching (PHA-L), core fucosylation (LCA), mannosylation (ConA), or α2,3-linked sialylation (MAL-1) of the bulk plasma glycoproteins (Fig. 1C). This suggests that FcRn-mediated deficiencies in sialylation are specific to IgG, which is consistent with reports showing that FcRn-mediated recycling is specific to IgG and the non-glycosylated plasma protein albumin (*24, 31*).

Additionally, we stained tissue sections with SNA to confirm that FcRn-mediated defects in sialylation were not observed at the tissue level. There were no observable (Fig. 1D) or statistical (Fig. 1E) differences in the fluorescence intensity of SNA between wild type and FcRnKO animals in lung, liver, spleen, or kidney. Together, these data show that FcRnKO animals exhibit a selective IgG sialylation deficit that is not observed in tissue or bulk plasma glycoproteins.

### Endothelial cells are capable of IgG sialylation

We next asked whether endothelial cells, the primary cell thought to be responsible for IgG recycling, are capable of sialylating IgG *ex vivo*. To do this, we isolated CD31^+^ lung endothelial cells from WT and FcRnKO mice and confirmed by quantitative PCR that both strains express ST6Gal1 in culture (Fig. 2A), suggesting that they have the enzymatic machinery necessary for sialylation, but that only wild type and not FcRnKO cells express FcRn (Fig. 2B). Moreover, CD31^+^ cells from the lungs of both FcRnKO and WT mice showed a high and indistinguishable degree of surface α2,6-sialylation as measured by SNA flow cytometry (Fig. 2C-2D), confirming that ST6Gal1 is enzymatically active in both genotypes.

**Fig. 2.**
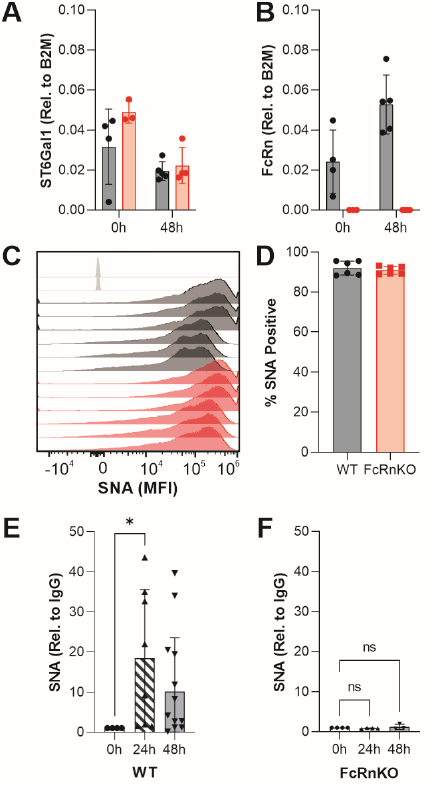
Primary murine *ex vivo* endothelial cells sialylate IgG in culture. Wild type and FcRnKO endothelial cells were isolated from lung tissue. Cells from both strain backgrounds express ST6Gal1 transcript following isolation from the animal (**A**), and only WT animals express FcRn (**B**). Cells from both strains have surface sialylation, as shown through flow cytometry (**C-D**). IgG was incubated with primary endothelial cells for 24 and 48 hours, then re-isolated from the media. Sialylation status was assessed via SNA ELISA. IgG was significantly more sialylated following incubation with WT (**E**) but not FcRnKO cells (**F**).

To test whether these cells were capable of IgG sialylation, we enzymatically removed the sialic acid from bulk IgG preparations to ensure available sialylation sites and incubated CD31^+^ primary lung endothelial cells with the enzymatically modified IgG. After 24 or 48 hours, IgG was purified from the conditioned media. We found that the sialylation status had increased approximately 13-fold over the input IgG by wild type cells (Fig. 2E); however, IgG recovered from the media of FcRnKO endothelial cells did not exhibit the same increase in sialylation (Fig. 2F). These findings closely align with endogenous IgG sialylation isolated directly from wild type and FcRnKO animals (Fig. 1A) and collectively indicate that not only is FcRn critical for IgG sialylation, but that endothelial cells specifically are capable of IgG endocytosis, addition of sialic acid with an α2,6 linkage, and subsequent IgG release back into the local environment.

It is accepted, however, that endothelial cells are not the only cells that express FcRn. Cells of hematopoietic origin have also been demonstrated to express FcRn, and some studies have suggested that FcRn^+^ hematopoietic cells may also participate in IgG recycling, although the extent to which they do so remains unclear (*23, 28*). Thus, we derived macrophages from murine bone marrow (BMDMs) and polarized the cells towards an M1 or M2 phenotype with LPS and IFNγ or IL-4 respectively. We found that ST6Gal1 was expressed at a slightly lower quantity than in primary endothelial cells and that it was expressed more strongly in resting M0 and M2 macrophages relative to M1 macrophages, regardless of the strain background (Fig. 3A). Similarly, FcRn was also present in wild type M0 and M2 macrophages, and FcRnKO macrophages lacked FcRn (Fig. 3B). Mirroring ST6Gal1 expression, wild type M1 macrophages significantly downregulated FcRn. Cells of both genotypes and under all polarization conditions were similarly α2,6 sialylated at the cell surface, as measured by SNA flow cytometry (Fig. 3C-E).

**Figure 3.**
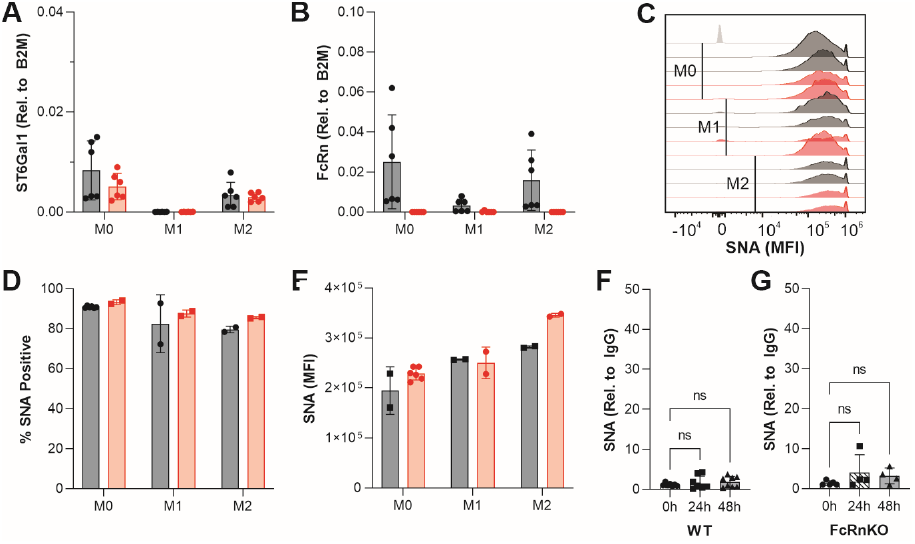
Bone marrow-derived murine macrophages do not sialylate IgG in culture. Wild type and FcRnKO macrophages were derived from bone marrow, then polarized towards M1 or M2 phenotype. Non-polarized (M0) and M2 macrophages expressed ST6Gal1 transcript in both strain backgrounds, while only WT macrophages expressed FcRn (**A-B**). M1 macrophages expressed low levels of ST6Gal1 and FcRn regardless of strain background. However, macrophages from all polarization conditions and strains were highly SNA positive by flow cytometry (**C**). No difference was noted in the percentage of cells that were SNA^+^ (**D**), or in the MFI of SNA signal (**E**). Cells were then isolated with IgG, and IgG was re-isolated after 24 or 48 hours as described with endothelial cells. In contrast to primary lung endothelial cells, macrophages from both conditions failed to robustly sialylate IgG (**F-G**).

Based on the ST6Gal1 and FcRn expression data, we incubated desialylated IgG with resting M0 macrophages for 24 or 48 hours. Unlike with primary endothelial cells, IgG isolated from the conditioned media showed no significant change in sialylation even after 48 hours (Fig. 3F-3G). Moreover, this lack of change was seen in both wild type and FcRnKO backgrounds. These data indicate that although FcRn is necessary for IgG sialylation *in vitro* and *in vivo*, the concomitant expression of FcRn and ST6Gal1 is insufficient for the cell to be involved in IgG sialylation.

### IgG colocalizes with FcRn and ST6Gal1 inside of human endothelial cells

Having identified murine endothelial cells as a potential site of IgG sialylation, we next sought to confirm these findings in a human system using the HMEC-1 endothelial cell line. We first confirmed the expression of several glycosyltransferases, including ST6Gal1, in HMEC-1 cells (Fig. 4A). The cells express ST6Gal1 but not ST6Gal2, which is reportedly expressed only in the central nervous system. Importantly, flow cytometry showed that HMEC-1 cells not only carried α2,6 sialylation at the cell surface (Fig. 4B), but they also robustly expressed FcRn (Fig. 4C), confirming that the glycosylation and receptor profile of these cells were similar to primary murine lung endothelial cells.

**Figure 4.**
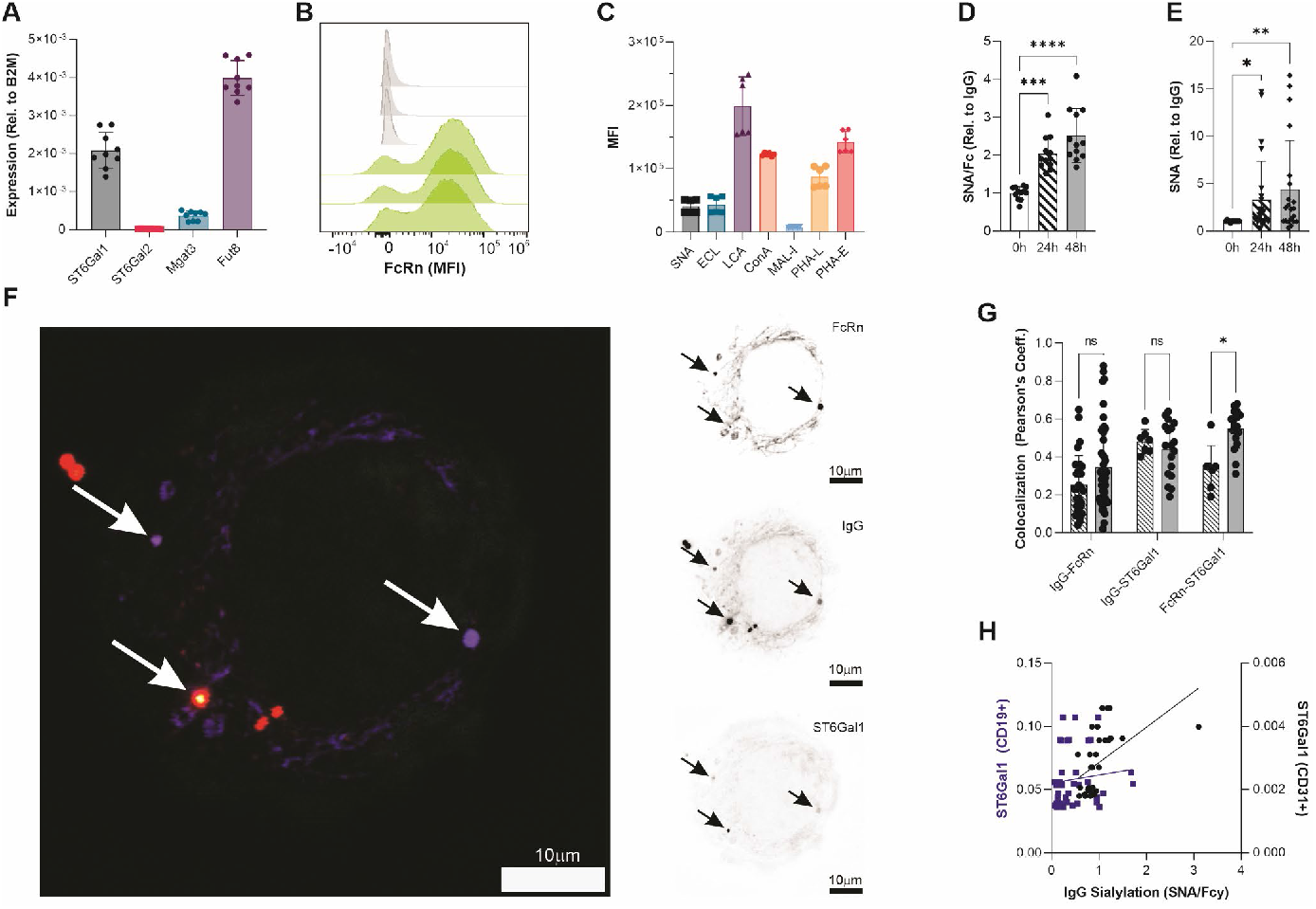
Human endothelial cells are capable of IgG sialylation. The HMEC-1 cell line expresses sufficient transcript levels of the glycosyltransferases ST6Gal1, Mgat3, and Fut8, but not ST6Gal2 (**A**). By flow cytometry, HMEC-1 cells express FcRn (**B**) and normal levels of surface glycosylation (**C**). Following 24 and 48-hour incubation with HMEC-1 cells, the sialylation status of IgG recovered from the media was significantly increased (**D**). The sialylation status of biotinylated IgG captured from HMEC-1 media using streptavidin was likewise increased (**E**). Confocal imaging showed that IgG co-localizes with ST6Gal1 and FcRn intracellularly after 24 and 48 hours of incubation, as illustrated by a representative image following 48-hour incubation (**F**). Points of co-localization are demarcated by white arrows. Individual color channels were inverted for clarity to show the localization of FcRn, IgG, and ST6Gal1 independently. Co-localization of each marker pair, as indicated, was quantified using ImageJ across many images (**G**). Mice were euthanized and ST6Gal1 expression from splenic CD19^+^ B cells and CD31^+^ lung endothelial cells was correlated with the sialylation status of plasma IgG, demonstrating that endothelial cell ST6Gal1 more strongly correlates with IgG sialylation *in vivo* than B cell ST6Gal1 (**H;** n > 33).

Given that HMEC-1 cells possessed what we believe to be the necessary components of the reaction, we next quantified the ability of these cells to add sialic acid to exogenous IgG. As before, we incubated HMEC-1 cells with IgG for 24 and 48 hours. IgG purified from the conditioned media carried a significant increase in IgG sialylation (Fig. 4D), supporting our hypothesis that both murine and human endothelial cells are responsible for IgG sialylation.

To further confirm that all signal stemming from the isolated IgG was solely due to the added human IgG, we also incubated HMEC-1 cells with purified biotinylated IgG. Using streptavidin capture rather than Protein A purification, we once again found a robust increase in IgG sialylation, as detected by SNA (Fig. 4E). These data reveal that both murine and human endothelial cells are able to take up exogenous IgG, add α2,6-linked sialic acid, and re-release the sialylated form of IgG back into the local environment.

In our previous work, we found that B cells are inefficient at IgG sialylation due to intracellular trafficking that limits ST6Gal1 and IgG co-localization (*20*). We therefore reasoned that in order for endothelial cells to sialylate IgG, ST6Gal1 must be able to contact ingested IgG within the intracellular environment. Having shown that HMEC-1 cells are capable of IgG sialylation (Fig. 4D-4E), we incubated them with IgG for 24 or 48 hours and examined the colocalization between IgG, FcRn, and ST6Gal1 by confocal microscopy. When we quantified two-color co-localization across all images taken, we observed no change in the co-localization between IgG and FcRn between 24 and 48 hours (Fig. 4F). However, the amount of co-localization between FcRn and ST6Gal1, as well as the co-localization between IgG and ST6Gal1 increased between 24 and 48 hours (Fig. 4D), suggesting that ST6Gal1 is being increasingly trafficked from the *trans*-Golgi to the endo/lysosomes containing both IgG and FcRn. Further, in the representative images shown from the 48-hour incubation, we observed distinct points where IgG, FcRn, and ST6Gal1 signal coalesce in a single vesicle (Fig. 4G). These data confirm that ST6Gal1 has direct access to the endo/lysosomal compartments known to carry both ingested IgG and FcRn within human endothelial cells, and contrasts with what we observed in antibody-secreting B cells which poorly sialylate IgG (*20*).

Data obtained from *in vitro* and *ex vivo* model systems predict that ST6Gal1 expression in endothelial cells should more accurately reflect overall IgG sialylation than ST6Gal1 expression within the B cell compartment. To test this hypothesis in an *in vivo* system, we immunized mice to boost their IgG titers and to stimulate an immune response, isolated both CD19^+^ B cells and CD31^+^ endothelial cells, and purified RNA for quantitative PCR analysis. As the model predicts, the sialylation status of IgG from the plasma of the same animal strongly correlated with the expression of ST6Gal1 in CD31^+^ endothelial cells but poorly correlated with ST6Gal1 in CD19^+^ B cells (Fig. 4H).

Collectively, these data support a novel paradigm in which the FcRn-mediated IgG recycling pathway in endothelial cells is expanded to include IgG-specific glycan remodeling as a means to dynamically alter both the circulatory lifetime and functional activity of endogenous as well as exogenous IgG and potentially other Fc domain-based therapeutics. The discovery of this pathway thereby opens exciting possibilities for novel treatments aimed at manipulating endogenous IgG to address a variety of inflammatory and autoimmune conditions.

## Acknowledgments

We would like to thank Jill Cavanaugh for sample preparation, particularly in performing mouse necropsies. We also graciously thank Dr. James C. Paulson (Scripps) for providing ST6Gal1KO samples.

## Author contributions

Conceptualization: LMG, BAC

Methodology: LMG, BAC

Investigation: LMG, KMR, ENK, MDL, BAC

Formal analysis: LMG

Visualization: LMG, BAC

Funding acquisition: BAC

Project administration: LMG, BAC

Supervision: BAC

Writing – original draft: LMG

Writing – review & editing: LMG, BAC

## Data and materials availability

All data are available in the main text or the supplementary materials.

## Materials and Methods

### Endothelial cell harvest, derivation of murine macrophages, and IgG isolation from plasma

Mice were sacrificed by CO_2_ inhalation according to a CWRU-approved IACUC protocol, and blood was obtained via cardiac blood draw into 3.2% sodium citrate to prevent coagulation. Plasma was generated from blood by taking the liquid fraction following centrifugation at 2500 x g for 15 minutes. IgG was isolated from murine plasma via Protein A affinity chromatography, and buffer exchanged into PBS for further analysis.

Endothelial cells were obtained by mincing the lung with scissors followed by digestion with 3 mg/mL collagenase I. Cells were filtered through a 70 μm strainer, and all CD31^+^ cells from one lung were plated into 3 wells of a 12-well plate in DMEM (see below for description of complete media). Adherent cells were rinsed gently with PBS daily for three days, and experiments were carried out once cells reached confluency, which was approximately 3 days.

Murine femurs were harvested and cut at one end, and bone marrow was spun out by centrifugation at 2,000 x g for 10 min. Bone marrow cells were resuspended in complete BMDM media (see “Cell culture” for description of complete media) and 5 million cells were plated in 100 mm plates. Bone marrow-derived macrophages (BMDMs) were supplemented with an additional 2 mL of BMDM media and 1 mL of L929 conditioned media on day 4, and on day 7, cells were re-plated prior to polarization or usage in experiments. Polarization to M1 was carried out via 24-hour stimulation with 5 ng/mL LPS and 50 ng/mL IFNγ. Polarization to M2 was carried out via 24-hour stimulation with 20 ng/mL IL-4.

### Tissue histology & confocal microscopy

Tissues were harvested following sacrifice via CO_2_ inhalation and fixed in 10 % formalin (VWR, Radnor, PA) for 24 hours. Tissues were sent to AML Laboratories (Jacksonville, FL) to be embedded onto slides and stained with H&E. Unstained sections were returned for immunofluorescence staining with 5 μg/mL SNA-FITC (Vector Labs). Images of H&E-stained sections were acquired using a High-Speed Microscope Camera (Amscope, Irvine, CA) on a Leica TCS SP5 confocal microscope (Leica, Wetzlar, Germany). Images of SNA-stained sections were acquired via the Leica TCS SP5 confocal microscope (Leica, Wetzlar, Germany).

### Cell culture

#### BMDM media

Advanced RPMI (Gibco) supplemented with penicillin/streptomycin, L-glutamine, and 10 % fetal bovine serum. 25 % of the culture volume was made up of L929 conditioned media to supply M-CSF as a growth factor. Cells were used upon derivation without further passaging.

#### Primary lung endothelial cell media

Advanced DMEM (Gibco) supplemented with penicillin/streptomycin, L-glutamine, 20 % fetal bovine serum, 100 μg/mL heparin, and 50 μg/mL endothelial cell growth supplement. Cells were used upon isolation without passaging.

#### HMEC-1 media

MCDB131 (Gibco) supplemented with penicillin/streptomycin, L-glutamine, 10 % fetal bovine serum, 1 μg/mL hydrocortisone, and 10 ng/mL epidermal growth factor. To passage, media was removed, cells were washed with PBS, then trypsinized at 37 °C with 0.05 % trypsin-EDTA (Gibco 25300-054) until the cell layer was dispersed. Complete culture medium was added to inhibit trypsin, cells were pelleted at 300 x g for 5 minutes, and the pellet was resuspended in fresh media. Cells were used for experiments until passage 15.

All cells and cell lines were incubated at 37 °C and 5 % CO_2_. HMEC-1 cells were purchased from ATCC (CRL-3243) and utilized until passage 15 as described.

### IgG incubation assay

Sialic acids were removed from human polyclonal IgG (Innovative Research IHIUGGAP) via recombinant neuraminidase (NEB P0720L) as described by the manufacturer. In brief, 1 mg of protein was incubated at 37 °C with enzyme and buffer for at least 16 hours to allow the reaction to reach completion. 50 μg of the completed reaction was then added to the media of cells plated in a 6-well plate. Media from control wells was immediately harvested, and input IgG re-isolated from the media as described elsewhere.

### Image analysis and quantification

Images were quantified using ImageJ software to determine the mean pixel intensity of each lectin on a per-image basis. ImageJ was also used to determine degree of two-color colocalization via the Pearson’s coefficient. Images of one cell at a time were taken wherever possible to determine degree of colocalization.

### Lectin ELISA and streptavidin capture

Purified IgG was diluted to 1 ng/μL in PBS, pipetted into a 96-well high-binding ELISA plate (Microlon High Binding; Greiner BioOne), and incubated overnight at 4 °C. The plate was blocked with carbohydrate free blocking solution (Vector Labs) for 1 hour at room temperature. Biotinylated lectins (Vector Labs) were diluted to 1 μg/mL in carbohydrate free blocking solution and incubated on the plate for 1 hour at room temperature.

Signal was detected using europium-conjugated streptavidin (Perkin Elmer) and time-resolved fluorescence as measured in a Victor Nivo plate reader. Alternatively, 100 μL of cell culture media was pipetted into a 96-well Immobilizer Streptavidin plate (ThermoFisher 436014) and incubated overnight at 4 °C. Plates were washed, and FITC-conjugated lectins (Vector Labs) diluted to 1 μg/mL or AF488-conjugated anti-human-IgG antibody (Jackson 309-065-008) diluted 1:500 in carbohydrate free blocking solution were incubated overnight. Plates were washed, 100 μL of PBS was pipetted into each well, and fluorescence was read using a Victor Nivo plate reader.

### Flow cytometry

Bone marrow-derived macrophages and primary endothelial cells were isolated as described above. HMEC-1 cells were lifted as described for passaging above and resuspended in carbohydrate-free blocking solution (Vector Labs) 30 minutes with Fc block. Cells were stained with SNA-FITC (Vector Labs) and FcRn-AF647 (sc-393064 AF647). Flow cytometry was run on a Attune NxT flow cytometer and data were analyzed using FlowJo.

### Quantitative PCR

RNA transcripts were isolated from cells using a Qiagen RNEasy Mini RNA isolation kit (Qiagen) per the manufacturer’s instructions. RNA was then converted to cDNA using a RevertAid First Strand cDNA synthesis kit (ThermoFisher). 1.5 μg of cDNA was then plated for quantitative PCR with Taqman Fast Advanced Master Mix (ThermoFisher). ΔCT relative to β2M was then visualized via GraphPad Prism as described below.

#### Primers used (Taqman)

Murine ST6Gal1: Mm01175607_m1

Murine FcRn: Mm01205449_g1

Murine β2-microglobulin: Mm00437762_m1

Human ST6Gal1: Hs00949382_m1

Human FcRn: Hs00175415_m1

Human β2-microglobulin: Hs00187842_m1

### Data & statistical analysis

All data were analyzed and plotted using GraphPad Prism. Statistical analyses used were Student’s t-test or one-way ANOVA where appropriate (*p < 0.05; **p < 0.01; ***p < 0.001; ****p < 0.0001)

